# EHMT2/G9a-Inhibition Reprograms Cancer-Associated Fibroblasts (CAFs) to a More Differentiated, Less Proliferative and Invasive State

**DOI:** 10.1101/2023.07.17.549300

**Authors:** Nila C Wu, Rene Quevedo, Michelle Nurse, Kebria Hezaveh, Haijiao Liu, Fumao Sun, Julien Muffat, Yu Sun, Craig A Simmons, Tracy L McGaha, Panagiotis Prinos, Cheryl H Arrowsmith, Laurie Ailles, Elisa D’Arcangelo, Alison P McGuigan

## Abstract

Cancer-associated fibroblasts (CAFs) have previously been shown to play a pivotal role in multiple cancer dynamics, including mediating tumor cell invasion: their pro-invasive secretory profile and ability to remodel the extracellular matrix (ECM) architecture particularly promote tumor progression through tumor cell invasion into surrounding tissue areas and beyond. Given that reduced CAF abundance in tumors correlates with improved outcomes in various cancers, we set out to identify epigenetic targets involved in CAF activation in the tumor-stromal margin to reduce overall tumor aggressiveness. Using the GLAnCE (Gels for Live Analysis of Compartmentalized Environments) co-culture platform, we performed an image-based, phenotypic screen and identified EHMT2 (also known as G9a), an epigenetic enzyme that targets the methylation of histone 3 lysine 9 (H3K9), as the most potent modulator of CAF abundance and CAF-mediated tumor cell invasion. Transcriptomic and functional analysis of EHMT2-inhibited CAFs revealed the involvement of EHMT2 in driving CAFs towards a pro-invasive phenotype. Further, EHMT2 signaling mediated CAF hyperproliferation, a feature that is typically associated with activated fibroblasts present in tumors, but the molecular basis for which has not thus far been identified. This study suggests a role for EHMT2 as a regulator of CAF hyperproliferation within the tumor mass, which in turn magnifies CAF-induced pro-invasive effects on tumor cells.

## Introduction

Cancer-associated fibroblasts (CAFs) are one of the most abundant stromal cell fractions in the tumor microenvironment (TME) in many tumor types(*1–3*) and CAF abundance is associated with poorer clinical prognosis for many cancers(*4, 5*), including head and neck squamous cell carcinoma (HNSCC)(*6, 7*). Most CAFs arise from quiescent fibroblasts residing in normal tissue that are primed to become activated during wound-healing and inflammation(*8–10*). During tumorigenesis, chronic and sustained fibroblast activation occurs(*11*), due to signaling from tumor and immune cells, as well as other microenvironmental changes, which drives the fibroblasts into various sub-types of CAFs including myofibroblast-like (myCAF) that exhibit increased contractility(*12*), increased ECM-remodeling capacity(*13–15*), and increased secretion of various signaling molecules(*16*). Bidirectional signaling between CAFs and tumor cells actively recruits CAFs to infiltrate the tumor mass where they proliferate to further increase CAF abundance and CAF-tumor cell interactions within the TME. This process of CAF infiltration into the tumor mass generates an invasive zone at the tumor margin where disease progression is accelerated via CAF-induced tumor cell invasiveness mediated by direct CAF-tumor cell contact(*17*), establishment of CAF-generated ECM tracks for tumor cell dissemination(*14*), secretion of factors that promote epithelial to mesenchymal transition (EMT) in the tumor cell(*18–20*), altering the immune contexture(*21*), and production of proliferative factors that drive tumor cell growth(*22–25*). Previous work suggests that myCAFs specifically strongly impact tumor cell invasion(*17, 26–28*) and immunosuppression(*2, 29–31*). Given the abundance of CAFs in the TME and their various contributions to disease progression, combined with their relative genetic stability compared to tumor cells(*32, 33*), strategies to target and modulate CAF behavior are potentially useful therapeutically(*34*).

Previous attempts to target CAFs to reduce their contribution to tumor progression have had relatively limited success thus far due to a lack of CAF-specific surface markers, and CAF population heterogeneity(*34*). Approaches that target CAF state to either reduce CAF abundance(*35, 36*) or the extent of CAF activation (either by reducing activation or reducing the proportion of activated CAF sub-types in the tumor(*37, 38*) have reached Phase II clinical trials, but have failed to meet minimum efficacy requirements for clinical continuation(*9, 37, 39*). A better understanding of the molecular mechanisms regulating CAF state would provide an opportunity to expand on these approaches.

Given that genetic alterations in CAFs are extremely rare, the perpetually activated CAF phenotype may be regulated intrinsically at the epigenetic level(*40, 41*) and therefore epigenetic targeting to reprogram CAF phenotype is an emerging therapeutic strategy. For example, acetylation of STAT3 by p300 leads to a loss of expression of SHP-1 tyrosine phosphatase, which leads to sustained pro-invasive CAF activity, and therefore blocking these pathways (JAK/STAT3 and silencing of SHP-1) can revert the pro-invasive CAFs *in vitro* and *in vivo*(*42*). Furthermore, upcoming, and promising work modulating epigenetic targets for anticancer treatment(*43, 44*) and fibrotic diseases(*45*), provide further examples of the plausibility of altering CAF epigenetics as a therapeutic strategy. While epigenetic regulation of myofibroblasts, in particular in the context of fibrosis, has been the focus of intense study(*46–49*), our understanding of these mechanisms in the context of the tumor invasive margin remains incomplete, limiting the potential of targeting CAF function at this site.

Exploring the epigenetic regulation of human CAF states within the invasive margin, as it pertains to CAF recruitment and subsequent CAF-mediated changes in tumor cell properties, is challenging using standard model systems: in mouse models, human CAFs co-injected with tumor cells are rapidly replaced by murine CAFs(*50*), while conventional *in vitro* co-culture spheroid assays lack the spatial architecture and resolution needed for effectively monitoring both CAF recruitment into the invasive margin and CAF-induced changes in tumor cell phenotype in this zone(*51*). We previously reported a 3D *in vitro* culture platform called GLAnCE (Gels for Live Analysis of Compartmentalized Environments), that provides a model of the invasive tumor margin and allows spatial and temporal monitoring of cell-cell interactions using high-content imaging(*52*). Here we set out to use GLAnCE to screen and identify epigenetic enzymes that regulate the abundance of HNSCC CAFs in the invasive margin, as well as their properties that lead to CAF-induced tumor cell invasion. We found that inhibition of the epigenetic methyltransferase EHMT2 (also known as G9a) significantly reduced CAF abundance and CAF-induced invasive tumor strand formation in the tumor-stroma invasive margin. Transcriptomic profiling implicated EHMT2 as a key regulator of CAF hyperproliferation and commitment to the pro-invasive CAF phenotype. Our results suggest that EHMT2 suppression in HNSCC CAFs inhibits a pro-tumorigenic CAF phenotype, which impacts cancer cell invasiveness within the invasive margin. Further, our study showcases the use of an *in vitro,* fully human culture assay to model complex human CAF behaviours and identify regulators of human CAF state that accelerate tumor progression.

## Results

### GLAnCE enables quantification of CAF infiltration into the tumor compartment and CAF-induced changes in tumor invasiveness

We previously described an engineered tissue model of the tumor-stroma invasive margin, using the GLAnCE platform(*52*), that recapitulates CAF-induced changes in cancer cell phenotype. In GLAnCE, cancer cells and fibroblasts are cultured in a 3D-extracellular matrix (ECM) and micro-molded to create a 24-well array of thin, patterned gels in which each cell type occupies a distinct compartment (Figure 1Ai). Within these gels, cancer cells and fibroblasts interact initially along the single interface that delineates the adjacent, but contiguous hydrogel sections mimicking the tumor-stroma boundary (Figure 1Aii)(*52*). Over time, cancer and fibroblast cells mix across this interface, causing interface breakdown and the formation of an invasive region (invasive margin) containing both cell types. To facilitate real-time monitoring of these dynamics, we seed GLAnCE gels using tumor and/or CAF cells that stably express fluorescently-tagged proteins.

Using this system, we previously showed that co-culture of the head and neck squamous cell carcinoma (HNSCC) CAL33 cell line, with HNSCC patient-derived CAFs in GLAnCE induced CAF infiltration and CAF-dependent changes in cancer cell phenotype, reminiscent of the properties of the dispersive edge of carcinomas (Figure 1Aiii). Specifically, CAF-cancer cell co-cultures showed striking remodeling at the compartment interface over a 5-day timeframe, including growth and invasion of cancer cells into the CAF compartment (Figure 1Bi) and progressive widening of the invasive margin between the initially distinct cell compartments, as is seen *in vivo*(*69*). We also observed changes in tumor aggregate structure into invasive multi-cellular strands (Figure 1Bii). Importantly, we observed that cancer cells re-organized into invasive strand structures only at invasive margin and not in regions away from the mixing zone where direct CAF-cancer cell contact did not occur(*52*). Further, we previously reported that both soluble signaling and direct contact between CAFs and cancer cells in GLAnCE induced the transcription of various EMT-related genes in the cancer cells(*52*), consistent with previous reports demonstrating that pro-tumorigenic CAFs promote cancer cell invasion *in vivo* by inducing EMT(*16, 70*).

We first set out to better characterize the properties of the CAF invasive margin in GLAnCE, to confirm our culture model captured the expected changes associated with the invasive margin as observed *in vivo*(*71*). *In vivo*, ‘invasion-competent’ cancer cells release cytokine, chemokine and growth factor signals that activate and recruit CAFs, widening the invasive margin and increasing CAF-cancer cell contact and tumor-CAF crosstalk that regulates further cancer cell invasion(*26*). Similarly, CAFs in GLAnCE infiltrated the cancer cell compartment (Figure 1Ci). The extent of CAF infiltration in the invasive margin was quantified by measuring the increase in CAF abundance (indicated by GFP+ pixels) in a specified region of interest (ROI) adjacent to the initial compartment interface (Figure 1Cii). CAF infiltration was significantly greater when cancer cells were present in the adjacent compartment (comp. CC) compared to an acellular gel (comp. CAF MC), suggesting active signaling between the cancer cells and CAFs to enhance CAF recruitment (Figure 1Ciii-iv). Note that using these same images, we also quantified CAF abundance (using the mean gray value of the GFP+ CAFs) in a specified ROI in the CAF compartment located away from the invasive margin and as expected we observed increased CAF abundance in co-cultures, further suggesting that secreted factors from the tumor cells influenced CAF properties in GLAnCE, similar to those previously described *in vivo* (Figure 1Cv). The abundance of CAFs present in the invasive margin in GLAnCE cultures was sensitive to the number of CAFs initially seeded in the CAF compartment but appeared to plateau at very high initial seeding densities: GLAnCE cultures with 5K versus 10K CAFs resulted in a significant increase in CAF abundance in the invasive margin (Figure 1D), but increasing the initial CAF seeding density to 50K cells produced only a minimal, not statistically significant further increase in CAF occupied area. We speculate that at very high cell densities, infiltration of the CAFs into the invasive mixing zone becomes space limited and that CAF invasion, as measured by area occupied by CAFs in the ROI region, becomes a less sensitive metric of the inherent capacity of a CAF to invade into the tumor compartment. We therefore performed all next experiments at initial CAF seeding density of 10K CAFs.

**Figure 1.**
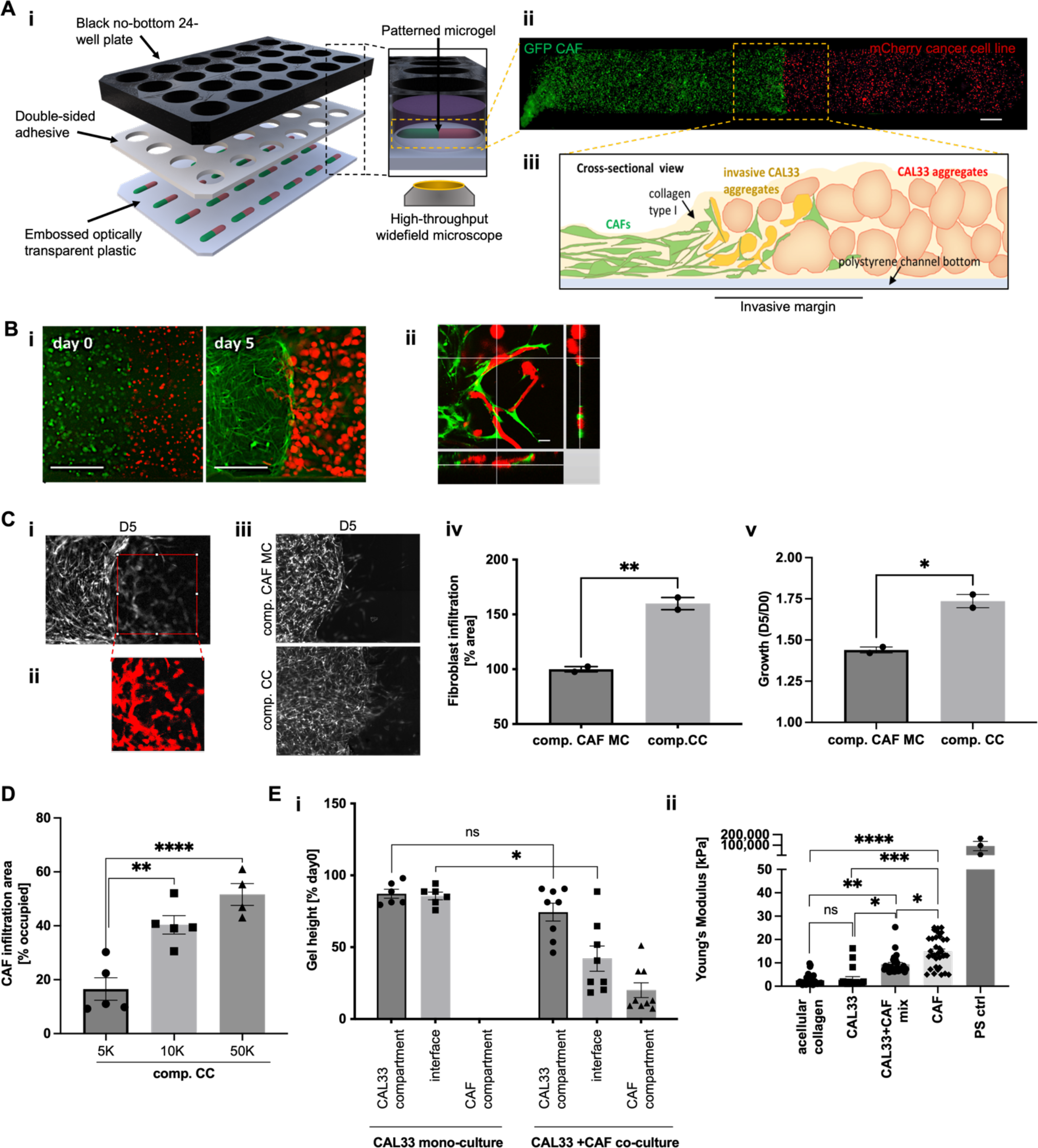
GLAnCE models the tumor-stroma invasive margin and captures CAF-induced changes in tumor invasiveness. **(A) (i)** GLAnCE assembly into a 24-well plate with standard dimensions and a custom-made optically transparent bottom that is seeded with cells embedded in a thin layer of hydrogel, with two adjacent compartments (patterned gel). Each well holds one compartmentalized gel, which can be imaged with an automated, widefield microscope. **(ii)** Image acquired of a full GLAnCE patterned gel where GFP CAFs are shown in green and mCherry CAL33s are shown in red. Scale bar is 500 µm. (iii) Cross-sectional schematic of spatially-defined cancer cell-CAF interface breakdown to generate tumor-stroma invasive margin, depicted by the yellow dotted selection in (Ai). **(B) (i)** Compartmentalized hydrogels after seeding (left) and at assay endpoint (day 5, right). CAFs are shown in green and CAL33 cancer cells in red; scale bar is 500 µm. **(ii)** Orthogonal view of cancer cells (red) arranging into invasive strands when in contact with CAFs (green) in the invasive margin; scale bar is 50 µm. **(C) (i)** Schematic of image analysis workflow to quantify CAF infiltration. A region-of-interest (ROI) is positioned adjacent to the CAF interface on day 0. **(ii)** The same ROI is positioned on the corresponding day 5 image, cropped and thresholded, and the % area of the mask is quantified as a measure of CAF infiltration. **(iii-iv)** Cancer cells recruit CAFs and enhance CAF infiltration into the invasive margin in co-cultures (comp. CC) compared to monocultures where CAFs migrate into an adjacent acellular gel (comp. CAF MC). Results show mean ± SEM for n = 2 for N = 2 biological replicates. **(v)** CAF growth in the CAF compartment away from the invasive margin is also influenced by the presence of an adjacent cancer cell population (comp. CC) compared to an adjacent acellular gel (comp. CAF MC). Results show mean ± SEM for n = 4 for N = 2 biological replicates. **(D)** CAF infiltration into the tumor compartment increases with increasing initial CAF seeding numbers. Results show mean ± SEM for N = 4/5 biological replicates. **(i)** Hydrogel contraction in the z-dimension increases in regions of GLAnCE cultures where CAFs are present. Results show mean ± SEM for N = 3 biological replicates. **(ii)** Hydrogel stiffness increases in regions of GLAnCE where CAFs are present. Results show mean ± SEM for N = 3 biological replicates.

Upon invasion *in vivo*, CAFs provide routes for invasive cancer cells to escape their ECM confinement by remodeling and stiffening of the ECM; this has also been observed in HNSCC(*6, 14, 72, 73*). Utilizing SEM, we identified prominent pores in ECM hydrogels remodelled by CAFs (Figure S1A). Specifically, ECM pores were only observed in hydrogel cultures containing CAFs (mono-cultures or CAF-cancer cell mixed co-cultures), but were completely absent in tumor-only hydrogel cultures and a-cellular hydrogels, suggesting that the ECM tracks were generated by the CAF cells. Given that cancer cells alone did not generate ECM pores in the timeline of our assay, we speculate that the invasive cancer cell strands observed in co-cultures take advantage of the CAF-generated ECM spaces, as other studies have reported(*14, 74, 75*). In GLAnCE co-cultures, regions where CAFs were present additionally exhibited hydrogel contraction (both in the CAF compartment and the invasive margin) to different thicknesses at endpoint (Figure 1Ei). As expected, the greatest contraction was observed in the CAF compartment, followed by an intermediate level in the invasive margin, and little to no hydrogel contraction in the cancer cell compartment, suggesting that CAFs primarily mediated hydrogel remodeling. Consistent with these observations, when we measured the stiffness of GLAnCE hydrogels directly using AFM, we found that as expected, hydrogels populated by CAF monocultures displayed the greatest change in stiffness compared to acellular hydrogels or cancer cell mono-cultures, while CAF-cancer cell mixed co-cultures had intermediate stiffness values (Figure 1Eii). Together, this data suggests that GLAnCE cultures recapitulate CAF-dependent characteristics of the tumor-stroma invasive margin. This comprises bulk ECM contraction, significant stiffening, and the generation of invasive tracks, which can then facilitate cancer cell invasion at the tumor-stroma interface. Further, cancer cells undergo phenotypic changes in the presence of CAFs, and this interaction is necessary to facilitate the formation of invasive tumor strands.

### Epigenetic modulation of CAFs impacts infiltration into the invasive margin and CAF-induced tumor invasiveness

Having validated our GLAnCE model of the invasive tumor margin, we next set out to identify epigenetic regulators of CAF infiltration into the invasive margin. Specifically, we aimed to identify targets that reduce CAF infiltration into the tumor compartment, thereby decreasing the size of the invasive margin and reducing the formation of invasive tumor cell strands. Because epigenetic machinery has previously been proposed to control fibroblast activation(*41, 46, 47, 76*), we assessed the impact of CAF treatment with chemical probes provided by the Structural Genomics Consortium (SGC), which target epigenetic enzymes. A library of 32 probes was utilized, with each probe being a potent and selective inhibitor with cellular activity at ≤1 µM(*77, 78*) (*SI Table 1*). Specifically, CAFs were seeded in 2D monoculture and treated with probes at a standard dose of 1 µM for 7 days, with one round of drug replenishing on day 5. Treated CAFs were then detached from 2D and counted, to seed an equal number of CAFs into each GLAnCE co-culture gel. After 5 days of compartmentalized co-culture in GLAnCE, hydrogels were assessed for CAF abundance within the invasive margin, as a measure of CAF infiltration, and number of tumor strand structures, as a measure of CAF-induced cancer cell invasiveness (Figure 2A).

**Figure 2.**
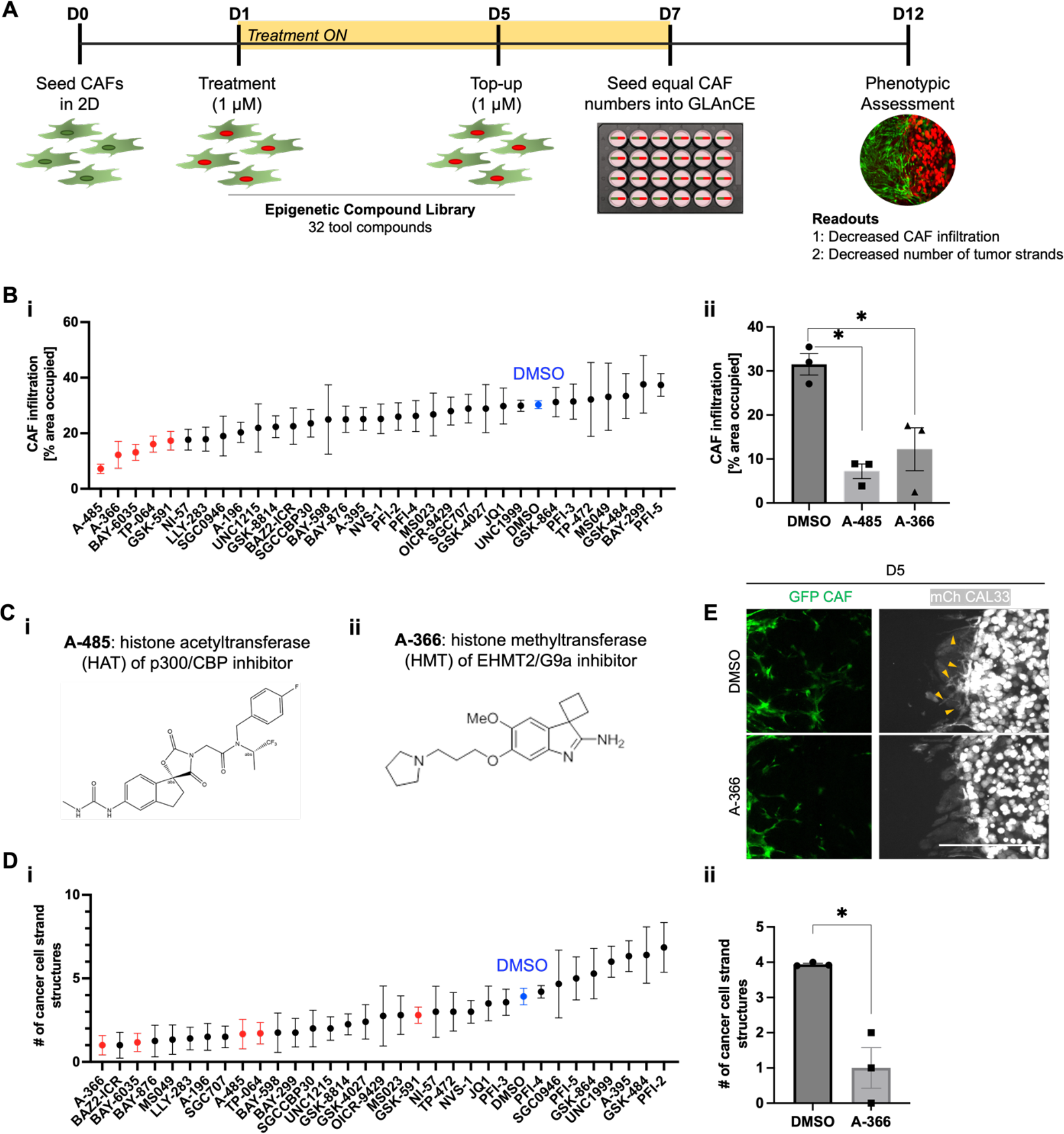
Epigenetic modulation of CAFs impacts CAF infiltration and CAF-induced cancer cell strand formation. **(A)** Experimental workflow of probe screen in which CAFs were treated with different epigenetic enzyme regulators and then the properties of CAFs in GLAnCE co-cultures was assessed. **(B) (i)** Effect of epigenetic probe library on CAF infiltration (readout 1), ranked by effect size. Top 5 compounds capable of reducing CAF infiltration are shown in red; DMSO control is shown in blue. Results show mean ± SEM for n = 3 biological replicates. **(ii)** Treatment of CAFs with probes A-485 and A-366A produced a significant reduction in CAF infiltration compared to DMSO control treatment. Results show mean ± SEM for n = 3 biological replicates. **(C)** Chemical structures of the tool compounds **(i)** A-485 (histone acetyltransferase of p300/CBP inhibitor) and **(ii)** A-366 (histone methyltransferase of EHMT2 inhibitor). **(D) (i)** Effect of epigenetic probe library on tumor strand formation (readout 2), ranked by effect size. Top 5 compounds capable of reducing CAF infiltration from (Bi) are shown in red; DMSO control is shown in blue. Results show mean ± SEM for n = 3 biological replicates. **(ii)** Treatment of CAFs with A-366 produced a significant reduction in tumor strand formation compared to DMSO control treatment. Results show mean ± SEM for n = 3 biological replicate **(E)** Representative interface images on day 5 of GLAnCE co-cultures, comparing control CAFs to A-366 treated CAFs. Yellow arrows indicate invasive strands, which are distinct 3D structures, as opposed to the cells growing in 2D at the GLAnCE plate bottom, as discussed in D’Arcangelo et al(*52*). Scale bar is 1 mm.

We first shortlisted probes that produced a significant reduction in CAF infiltration into the invasive margin (Figure 2Bi). The tool compound A-485, which targets histone acetyltransferase (HAT) p300, was the most effective at decreasing CAF infiltration. Given that p300 activity has previously been implicated in the maintenance of a pro-invasive CAF phenotype(*42*), p300 inhibition was expected to reduce CAF-infiltration and therefore provided confidence in the robustness of our infiltration screening metric. The second-top inhibitor of CAF infiltration was A-366, a small molecule inhibitor of euchromatic histone methyltransferase 2 (EHMT2), also known as G9a, that is responsible for methylating histone H3K9 in euchromatin(*79*). Both A-485 and A-366 significantly decreased CAF infiltration into the invasive margin zone by at least 50% compared to DMSO-treated CAFs (Figure 2Bii, 2C). We next shortlisted probes that resulted in a reduction in the number of tumor strand structures formed in the invasive margin, a measure of CAF-induced cancer cell invasiveness. Interestingly, we observed that compounds that decreased CAF infiltration did not always alter the number of tumor strands and vice versa (Figure 2Di). Despite this, A-366 was the top hit in inhibiting tumor strand formation. The top five compounds capable of inhibiting CAF infiltration were not all among the top inhibitors of tumor invasiveness, designated by the red dots in Figure 2Bi and 2Di. This discrepancy suggested that probe treatments could affect a variety of concurrent CAF properties, leading to either altered infiltration or altered CAF-tumor cell interaction, or both.

Given that we wanted to identify pathways that primarily reduced CAF infiltration and subsequently also produced an effect on tumor cell invasiveness (as measured by strand formation), we selected A-366 as an interesting candidate for follow-up experiments, as this compound was a top negative modulator of CAF infiltration and produced the greatest reduction in tumor strand structures (Figure 2Dii) as seen in Figure 2E. Moreover, A-366 targets EHMT2 (also known as G9a) which has previously been shown to influence fibroblast activation in the context of both lung and renal fibrosis(*49, 80*), and most recently was linked to TGF-β1-mediated chemokine expression in CAFs using mouse models(*81*). Further, G9a inhibition has been shown to impact cancer stemness in a variety of tumour types(*82*) and aggressiveness in breast cancer(*83*). Our data suggesting a pro-invasive role of CAFs with uninhibited G9a, suggests the possibility that CAFs may exert similar functions when G9a inhibition is carried out systemically *in vivo* and/or may further enhance tumour invasiveness in these settings. This may specifically be the case when assessing invasiveness of primary tumours, as well as the appearance of metastases(*84*). Global patient data analyses linking higher levels of G9a expression to reduced overall survival likely also capture expression of this protein within the stroma(*84, 85*).

### EHMT2-inhibition decreases both CAF infiltration and CAF-induced cancer cell invasiveness *in vitro*

Having identified A-366 as an interesting candidate for follow-up, we first assessed the cytotoxicity of this compound treatment at the 1 µM concentration used for the screen, to confirm that a reduction in CAF invasion was not simply due to CAF death in response to probe treatment. Upon staining with propidium iodide (PI), we observed no significant difference between CAFs with or without probe treatments, suggesting that A-366 treatment was not cytotoxic to CAFs, but rather in our GLAnCE invasion assay, probe treatment induced a phenotypic change in the CAF population resulting in the observed decrease in CAF infiltration (Figure 3A). Since CAFs were treated with epigenetic inhibitors prior to their addition to GLAnCE co-cultures, where they were then cultured in fresh probe-free media, we next assessed the extent to which A-366-treated CAFs showed inhibited target activity throughout the duration of the co-culture assay. To do this, we quantified the presence of histone marks H3K9me2 at different timepoints during probe treatment (days 2 and 6) and after probe removal (days 8 and 10), using immunofluorescent staining (Figure 3Bi-ii). A-366 treatment resulted in prolonged target repression over the entire timeframe associated with our GLAnCE invasion assay, even with removal of the probe compound from the culture media, as shown by the high percentage of H3K9 mark repression in probe-treated cells compared to untreated cells on days 8 and 10 (Figure 3Biii). We also assessed if greater repression could be achieved by increasing the probe treatment concentration to 5 µM, however we did not observe greater repression for A-366 at higher treatment doses, suggesting that the 1 µM treatment used in our screen was adequate to achieve maximum repression of enzyme activity in our *in vitro* assay (Figure 3Biv).

**Figure 3.**
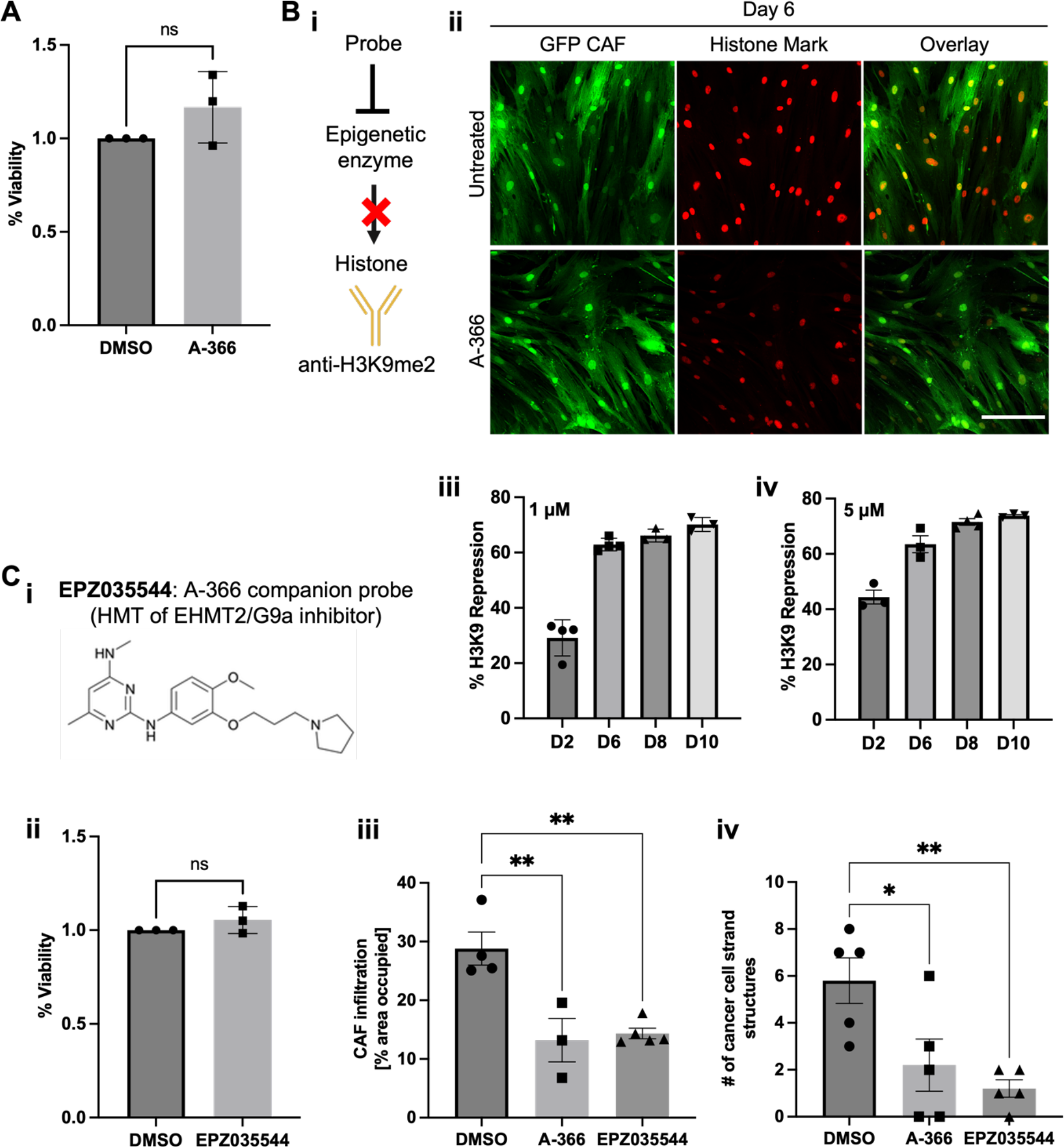
EHMT2 inhibition of CAFs is specific, prolonged, and robust. **(A)** Comparison of control and A-366 treatment on CAF viability. Cell viability staining (Propidium Iodide = PI indicates dead cells) of CAFs on day 7 of treatment indicated that A-366 treatment does not produce significant cell death of CAFs. Results show mean ± SEM for n = 3 biological replicates. **(B) (i)** Schematic representation of epigenetic modification and targeting of enzyme-dependent H3K9me2 histone mark to assess target repression. **(ii)** Representative 20X confocal immunofluorescence images probe-treated CAFs compared to DMSO-treated CAFs on day 6. Green indicates GFP CAFs and red indicates the histone mark (di-methylated H3K9). Scale bar is 200 µm. **(iii)** The histone probe mark remains repressed during 1µM A-366 treatment (day 2 and day 6), and also after the 1 µM A-366 treatment has been removed (day 8 and day 10), which corresponds to the time points during which CAFs were cultured in GLAnCE. Results show mean ± SEM for n = 3 biological replicates. **(iv)** Repression of the histone probe mark remained similarly repressed during and after a treatment with 5 µM of A-366, however treatment with the probe higher concentration did not increase histone repression compared to the 1 µM treatment. Results show mean ± SEM for n = 3 biological replicates. **(C) (i)** Schematic showing the chemical structure of the A-366 companion probe, EPZ035544, which also inhibits EHMT2. **(ii)** Dead cell staining (PI) of CAFs on day 7 of treatment with EPZ035544 indicated that the treatment does not produce significant cell death of CAFs. Results show mean ± SEM for n = 3 biological replicates. CAFs treated with the EPZ035544 compound similarly showed significantly reduced **(iii)** CAF infiltration and **(iv)** CAF-induced cancer cell strand formation. Results show mean ± SEM for n = 3 biological replicates.

We next confirmed that the inhibitory effects of A-366 on CAF infiltration were specific to the epigenetic enzyme EHMT2. To do this, we treated CAFs with a secondary companion compound, with the internal OICR identifier OICR0018881A01 and otherwise externally known as EPZ035544, which has a different molecular structure from A-366, but targets EHMT2 with similar potency and reported specificity(*86*) (Figure 3Ci). Treatment with this compound resulted in no significant toxic effects on CAFs (Figure 3Cii) and produced a reduction in CAF infiltration (Figure 3Ciii), and CAF-induced cancer cell invasive strand formation (Figure 3Civ) that was comparable to A-366 treatment. Overall, this suggested that inhibition of EHMT2 in CAFs was specific and long-lived, and led to a robust reduction in both CAF infiltration and CAF-induced tumor strand formation in the GLAnCE model of the tumor invasive margin.

Finally, to test whether EHMT2-inhibition in CAFs would produce the same phenotype when in GLAnCE cultures with another HNSCC cell line, we performed our GLAnCE invasion assay using CAL27 cells, another HNSSC cell line. As observed with CAL33 cancer cells, both A-366- and EPZ035544-treated CAFs co-cultured with CAL27 cells in GLAnCE exhibited decreased infiltration into the CAL27 tumor compartment, and induced less CAL27 invasive tumor strands, suggesting that EHMT2 inhibition produces a robust CAF phenotype in the context of multiple HNSSC cancer cell lines (Figure S1B). Taken together, our results suggest that modulation of the CAF epigenome using designed inhibitor probes results in changes in the *in vitro* infiltrative and pro-invasive properties of CAFs, and that inhibition of EHMT2 in CAFs specifically produced a significant decrease in both these properties.

### EHMT2-inhibition of CAFs produces transcriptional changes in fibroblast activation pathways

Next, to identify the molecular pathways that may be driving the phenotypic changes observed in CAFs upon EHMT2-inhibition, we performed transcriptomic profiling using bulk RNA-sequencing. RNA was isolated for RNA-seq from CAFs after 7 days of treatment with A-366, the companion probe EPZ035544, or vehicle (DMSO) (Figure 4A). In parallel, we also confirmed that the same treated batch of CAFs exhibited the expected phenotype in the GLAnCE invasion assay i.e. reduced CAF infiltration and reduced tumor strand formation (data not shown). Principal-component analysis (PCA) using Variance Stabilizing Transformation (VST) showed clustering of DMSO control samples versus probe-treated CAFs, with samples further clustered by the probe applied, although loosely (Figure 4B). Consistent with the fact that we targeted one enzyme specifically, we identified a relatively small group of genes (342 genes for A-366 vs DMSO, and 527 genes for EPZ035544 vs DMSO) that were significantly differentially expressed in probe treated CAFs compared to DMSO (false discovery rate (FDR) < 0.05) (Figure S1C). 146 differentially expressed genes (DEGs) were shared and upregulated between the two probe treatments, while 83 DEGs were shared and downregulated (35.7% combined); 46.6% and 17.6% of DEGs were exclusive to EPZ035544 and A-366 treatments, respectively (Figure 4C). We also compared the concordance of DEGs between the two different probe treatments by plotting the Log2FoldChange for the A-366 inhibitor against the EPZ035544 inhibitor and as expected, given that EHMT2 was the common target for both treatments, we observed excellent concordance of DEG directionality between treatments, with only 1/640 DEGs showing mismatched directionality.

**Figure 4.**
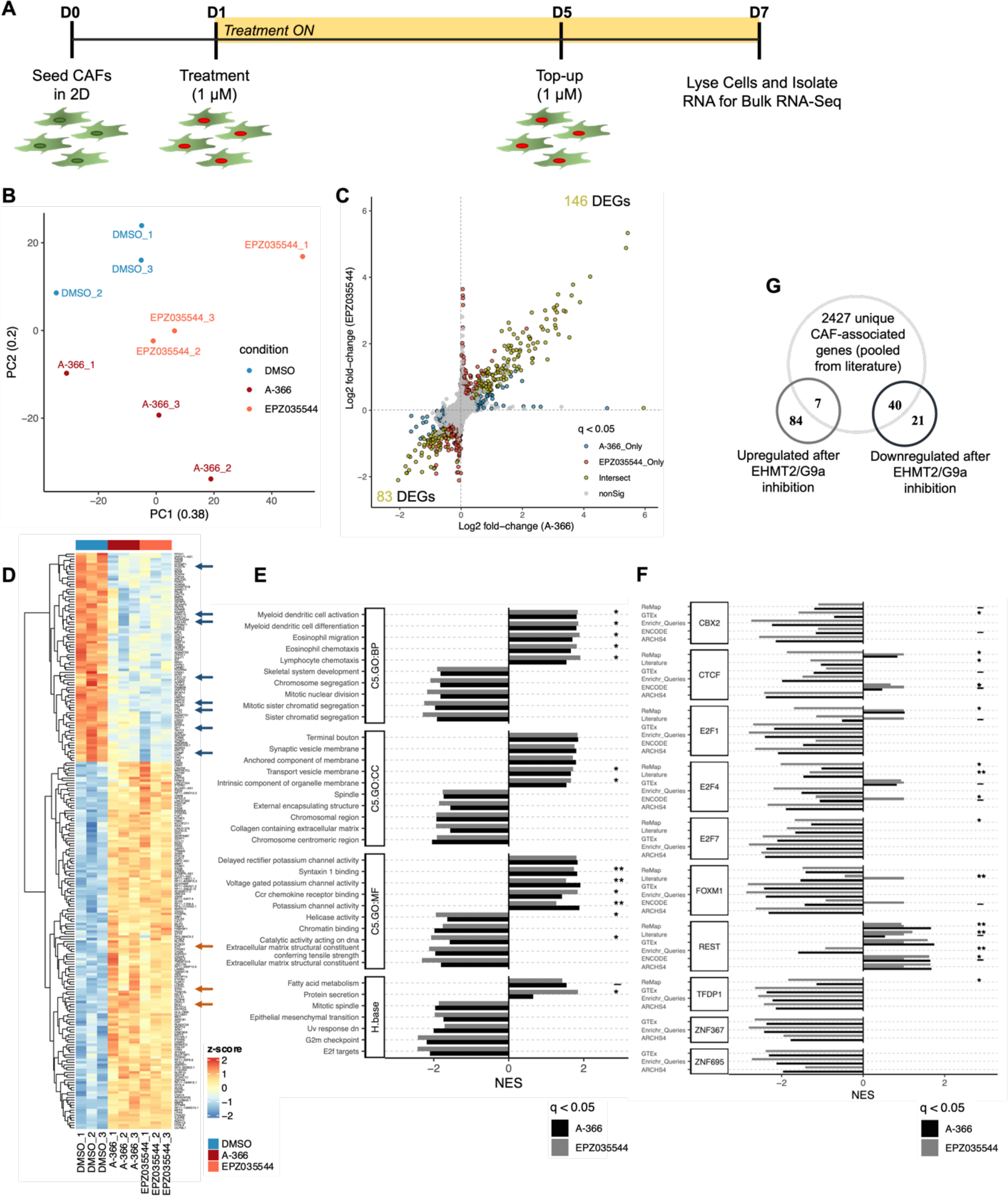
Transcriptomic characterization of EHMT2-inhibited CAFs by DEG, pathway and transcription factor activity analysis in our system. Our analysis revealed a shift away from the activated CAF phenotype towards a less pro-invasive state chiefly characterized by reduced expression of activation markers and pathway activity, reduced EMT-inducing capacity and upregulation of non-CAF pathways, coupled with a decreased proliferative capacity. **(A)** Schematic of bulk RNA-seq sample collection timepoint, of DMSO-, A-366, and EPZ035544-treated CAFs, 3 replicates per condition. **(B)** PCA plot of 9 samples, using Variance Stabilizing Transformation. **(C)** Plot of Log2FoldChange for the A-366 treatment vs. EPZ035544 treatment indicated good concordance of DEG directionality between treatments, with only 1/640 DEGs having mismatched directionality. **(D)** Heatmap for all overlapping 229 DEGs between A-366 and EPZ035544 treatment, compared to DMSO treatment. Several markers previously associated with the activated CAF phenotypes are present. **(E)** Visualization of the top 5 increased or decreased significant gene sets based on Fishers-p, quantified using the normalized enrichment score (NES) and grouped by the library used for analysis using the following libraries: Gene Ontology: Biological Pathways (GO:BP), Gene Ontology: Cellular Components (GO:CC), Gene Ontology: Molecular Functions (GO:MF), Hallmark collection (H.base). Annotations on the graph represent: gene set that is not significant in either condition alone but is significant when using the combined p-value (-), significant in EPZ035544-only (*), significant in A-366-only (**), and blank if significant in both EPZ035544 and A-366. **(F)** Transcription factor (TF) analysis using GSEA on gene sets defined by the ChEA3 tool, with the corresponding gene set libraries used for each significantly enriched TF listed on the y-axis. Graph annotations are the same as used in (E). Note that upregulation or downregulation profiles describe the targets associated with the TF, and not the TF expression levels itself. **(G)** Comparison of overlapping DEGs of probe-treated CAFs (EHMT2-inhibited CAFs) against a pooled CAF-associated marker repository(*94*).

Within the 229 overlapping DEGs between the two treatment groups (Figure 4D), we observed several genes that have previously been associated with fibroblast activation in the broad context of inflammation or disease, such as *LRRC15, COL5A2*, and *POSTN*(*87*). Further, some of these genes are overexpressed in HNSCC CAFs specifically, such as *LRRC15* and *COL5A2*, the expression of which has been associated with poor patient outcome and response to drug(*31*). Similarly, periostin (*POSTN)*, which was also downregulated in probe treated cells, has been associated with shorter overall survival of HNSCC patients(*88*) and has been shown to facilitate HNSCC cell-CAF invasion(*89*). Moreover, higher expression of *POSTN* in CAFs has been associated with an early activated CAF state that exhibited increased proliferation with higher expression, specifically at the tumor invasive margin(*88, 90*). Interestingly, we also observed the downregulation of genes associated with a universal fibroblast profile(*87*), such as *DPT, CXCL12* and *COMP* potentially suggesting a shift away from a fibroblast identity. Consistent with this idea, our upregulated overlapping DEG list contained genes that are highly expressed in neuronal progenitor cells and neurons that were not expected to be highly expressed in a fibroblast population, including: *BEX1*(*91*), *SYN-1*(*92*), and *SYP*(*93*). Further, when we compared probe-induced DEGs with a repository of CAF-associated gene markers defined from the literature across multiple CAF subtypes(*94*) (Figure 4G) we found relatively few upregulated CAF-associated DEGs (7/91) compared to downregulated CAF-associated DEGs (40/61), further suggesting a global reduction in the specification of an activated fibroblast state upon EHMT2 inhibition. Together, these observations suggest that the EHMT2-inhibited HNSCC CAFs showed gene changes previously associated with a diminished activated CAF phenotype.

We next set out to identify molecular pathways modulated by EHMT2 inhibition by performing gene set enrichment analysis (GSEA) between the two inhibitors and the control DMSO sample. We first confirmed that the significantly enriched gene sets between the two different inhibitor probes largely shared the same directionality (same NES) to increase the confidence of our interpretations (Figure S2A). In order to better distinguish between significant terms identified in either one or both treatments (EPZ035544 vs A-366), we qualified the top ranked gene sets, based on an FDR adjusted Fisher’s p-value, that were found to be significant in i) both treatments, ii) only in EPZ035544-treated CAFs (labelled *), iii) only in A-366-treated CAFs (labelled **), or only identified as statistically significant when both treatment datasets were combined (labelled -). We identified several pathways that aligned with our previous interpretation that EHMT2 inhibition decreases the level of CAF activation (Figure 4E). For example, we observed downregulation of ECM-remodeling pathways including ECM structural constituent and collagen containing ECM in probe treated cells, potentially suggesting that EHMT2 inhibition reduced CAF remodeling capabilities. EHMT2 inhibition also increased the expression of genes associated with fatty acid metabolism, a property of CAF progenitors as opposed to activated CAFs(*95*). Consistent with this, when we stained with BODIPY to evaluate lipid content, a measure of fatty acid metabolism, we observed that CAF treatment with the EPZ035544 probe produced a significant increase in BODIPY staining relative to DMSO-treated CAFs, while treatment with A-366 produced a similar but less robust (non-statistically significant) increase in BODIPY (Figure S2B). Probe treatment also resulted in downregulation of genes associated with the epithelial-to-mesenchymal transition (EMT) pathway including downregulation of a cohort of secreted signaling factors associated with EMT-induced cancer cell invasiveness including POSTN(*96*), CXCL12 (SDF1)(*97*), PTHLH(*98*), and PTX3(*99*). EHMT2 inhibition may therefore decrease the capacity of CAFs to elicit EMT in tumor cells through a decrease in paracrine signaling resulting in the observed reduction in CAF-induced tumor cell strand formation in the interface margin. Another notable set of pathways downregulated upon EHMT2 inhibition were associated with CAF proliferative capacity, including G2M checkpoint, E2F targets, and mitotic spindle pathways. Indeed, hyperproliferation has been reported as a key feature of the activated fibroblast phenotype(*100*); however, the epigenetic mediators regulating hyperproliferation in CAFs are not well understood(*9, 101, 102*).

Next, to determine if specific gene and pathway changes induced by EHMT2 inhibition were regulated by specific transcription factors, we performed transcription factor (TF) enrichment analysis using the ChEA3 tool(*66*), which comprises of several TF-target gene set libraries (Figure 4F). A relatively small number of TFs were found to be enriched: E2F1, E2F4, E2F7 and FOXM1 targets were significantly downregulated in probe-treated CAFs, while REST targets were significantly upregulated. The E2F transcription factor family plays an integral role in cell cycle progression and proliferation(*103*), and its downregulation aligns with our GSEA pathway analysis showing that EHMT2 inhibition produced decreased proliferation. FOXM1 has similarly been associated with increased cell proliferation(*104, 105*), and to be a driver of lung fibroblast activation in pulmonary fibrosis(*106*). Further, FOXM1 has been associated with regulating CAF-derived *COMP* secretion, which induces EMT in cancer cells(*107*), consistent with the decreased CAF-induced cancer cell strand formation observed in probe treated CAFs, where FOXM1 is downregulated. REST, also known as neuron-restrictive silencing factor (NRSF), is explicitly involved in the repression of neuronal genes in non-neuronal cells(*108*). Previous reports have shown that EHMT2 interacts with REST to effect repression of REST target genes, resulting in the inhibition of neuronal differentiation(*109, 110*). Upregulation of REST targets in our model may therefore be explained by EHMT2 inhibition preventing repression of REST target genes, thus reducing the repression of neuronal gene transcription programs. This is consistent with our observations that EHMT2 inhibition resulted in upregulated expression of neuronal genes *SYP* and *SYN-1* (Figure 4D) and increased staining for the neuronal marker *SYN-1*, a target gene of the REST transcription factor (Figure S2C). Reduced REST repression by EHMT2 inhibition may, therefore, relieve the silencing of competing (neuronal) transcriptomic programs, thereby altering the CAF activation state and consequent pro-invasive properties. This model of CAF reprogramming is indeed supported by previous observations of REST-mediated direct trans-differentiation of fibroblasts into neuronal progenitors and even mature neurons(*111*) where REST-silencing strategies allow for the expression of genes that specify terminal neuronal fate. Our data therefore suggests a deregulation of CAF phenotypic constraints upon EHMT2 inhibition in this assay.

### EHMT2-inhibition reduces CAF activation and curbs CAF hyperproliferation

Inhibition of EHMT2 resulted in reduced CAF abundance in the invasive margin and reduced CAF-mediated tumor invasive strand formation. Our transcriptional analysis suggested that the underlying mechanisms driving the changes in CAF behaviour upon EHMT2 inhibition resulted from a reduction in CAF activation predominantly via the downregulation of specific gene pathways that drive a CAF activation state (reduced EMT, reduced capacity for ECM organization, reduced expression of known CAF activation markers and reduced proliferative capacity), and via more subtle upregulation of pathways divergent from an activated state (increased FA metabolism and a potential shift towards a dedifferentiated state mediated by reduced repression of REST target genes). This altered CAF state appeared to be associated with changes in several CAF properties, some of which have previously been associated with EHMT2 signaling in fibroblasts(*49*) and other cell types(*112, 113*), and which may impact CAF abundance and CAF-induced tumor invasiveness in the invasive margin. We therefore set out to assess the impact of EHMT2/Ga inhibition on various CAF properties previously associated with an activated CAF state.

A recognized property of activated CAFs is increased contractility(*114, 115*), which can impact the ability to pull and re-organize matrix fibres during ECM remodeling(*116*), and generate intracellular traction forces to propel a cell forward during cell migration(*117*), both of which could potentially impact CAF infiltration into the invasive margin. We first assessed the impact of EHMT2 inhibition on CAF contractility using a 5-day gel contraction assay in which the extent of gel contraction reflects the capacity of cells to both pull on and reorganize the ECM fibres of the surrounding gel(*118*). Inhibition of EHMT2 produced no significant changes in gel contraction (Figure 5A) suggesting limited effects on CAF contractility. We then assessed the impact of EHMT2 inhibition on CAF migration in a Transwell® migration assay, in which the number of cells that migrate through the pores of the Transwell® membrane (no ECM is present) over a 24 hr period reflects the cells’ migratory capacity. In this assay, A-366 treatment resulted in a significant reduction in CAF migration, while EPZ035544-mediated inhibition produced a slight but not statistically significant reduction (Figure 5B). These observations suggest that, while EHMT2 inhibition in CAFs does not lead to robust changes in CAF contractility, it does result in lower CAF migration that may contribute to the observed reduction in CAF abundance in the invasive margin in our GLAnCE assay.

**Figure 5.**
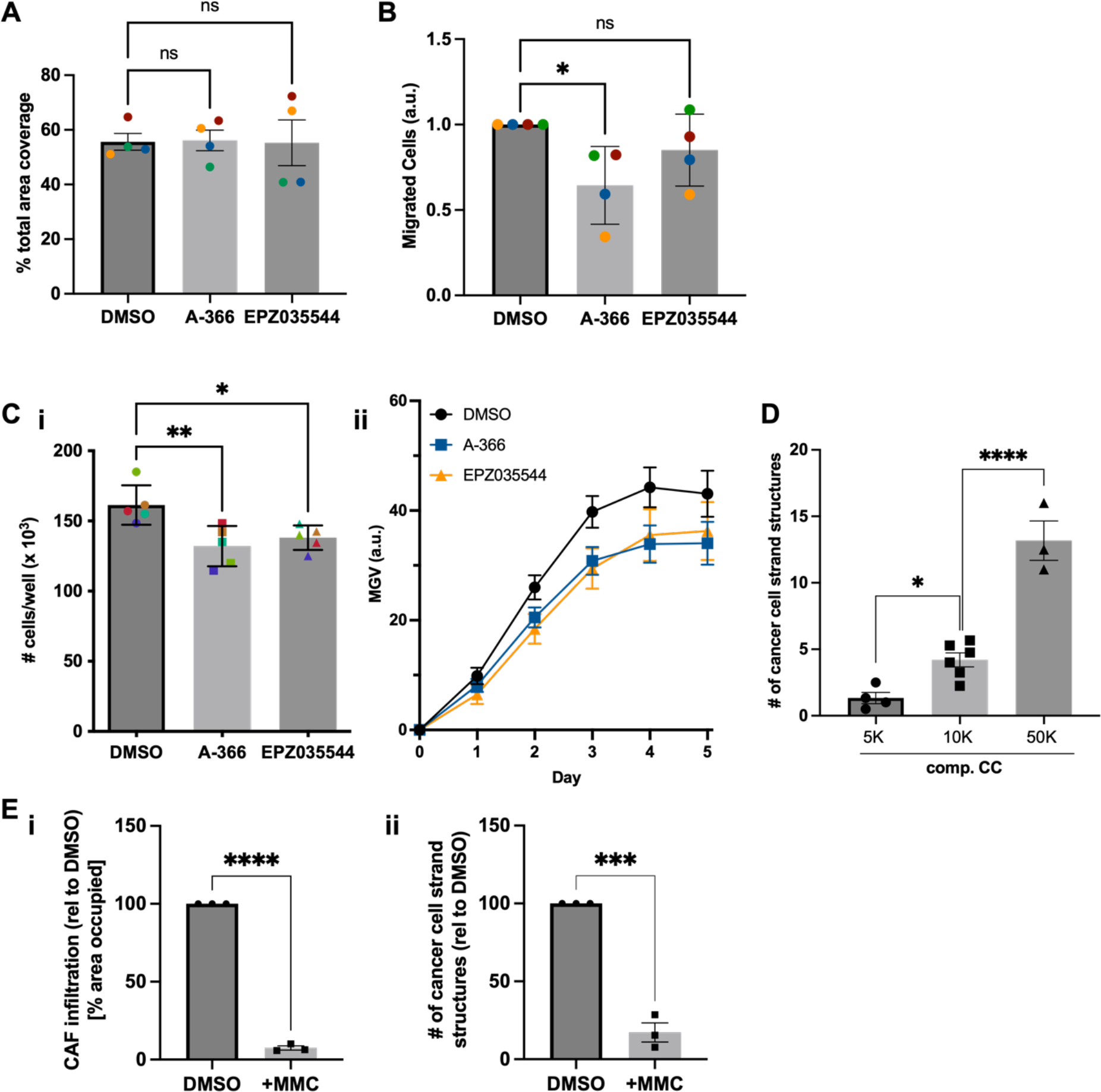
Functional assessment reveals that the decrease in tumor invasion is driven by alterations in an activated CAF phenotype upon EHMT2-inhibition. **(A)** Contraction assay performed by seeding gel plugs with CAFs treated with each condition, and gel area measured 5 days after seeding. Results show mean ± SEM for n = 4 biological replicates. **(B)** Migration assay performed using the Transwell® assay between treated CAFs, after allowing 24 hours for migration. Results show mean ± SEM for n = 4 biological replicates. **(C) (i)** Cell count of treated CAFs on day 7. Results show mean ± SEM for n = 5 biological replicates. **(ii)** Treated-CAF growth curves plotting change in GFP intensity. CAF growth is decreased upon EHMT2 inhibition. Results show mean ± SEM for n = 3 biological replicates. **(D)** Cancer cell strand number as a function of increasing CAF seeding numbers in compartmentalized GLAnCE co-cultures. The number of strand structures increase with increasing CAF supply in the adjacent compartment. Results show mean ± SEM for n = 3 biological replicates. **(E)** Inhibition of CAF proliferation with MMC (proliferation inhibitor) drastically decreased both **(i)** CAF infiltrative capacity and **(ii)** number of cancer cell strand structures. Results show mean ± SEM for n = 2 biological replicates, normalized to DMSO treatment of each replicate.

Another key feature of activated CAFs is hyperproliferation(*100*), the epigenetic mediators of which are not well understood. Given the 5-day timeline of the GLAnCE invasion assay, we hypothesized that CAF hyperproliferation could potentially contribute significantly to the abundance of CAFs within the invasive margin. EHMT2 inhibition using both probes produced a significant decrease in CAF number after the 7-day treatment regimen compared to DMSO-treated control CAFs (Figure 5Ci). Further, when we seeded treated vs untreated CAFs into GLAnCE to generate completely mixed co-cultures and monitored the growth of the CAF population using daily measurements of total mean gray value (MGV) of GFP intensity, we observed a persistent reduction in CAF abundance beyond the probe treatment period that is maintained over the period of the GLAnCE co-culture (5 days) (Figure 5Cii). These observations are consistent with previous studies that have shown that EHMT2 promotes proliferation during myogenic differentiation(*112, 113*). Altogether, this suggests that in addition to reduced cell migration, reduced proliferation of inhibited CAFs was a significant factor in reducing CAF abundance in the invasive margin.

In addition to reduced CAF abundance, tumor cell invasiveness was also reduced at the invasive margin in co-cultures with EHMT2-inhibited CAFs. To determine whether CAF abundance could by itself modulate the number of invasive tumor strand structures in our system and to further dissect the interplay between CAF proliferative rate and changes in CAF state suggested by our transcriptomics analysis, we assessed if an increased supply of CAFs was sufficient to increase tumor strand formation. When we increased the initial number of CAFs seeded in the CAF compartment, which led to increased CAF abundance in the invasive margin (Figure 1D), we observed a corresponding increase in the number of cancer cell strand structures (Figure 5D). This suggested that the capacity of tumor cells to form invasive strands was related to the abundance of CAFs within the tumor cell compartment and that treatments to reduce CAF availability could lead to reduced tumor cell invasiveness. Consistent with this, when we pre-treated CAFs with mitomycin C (MMC), an inhibitor of DNA synthesis that prevents cell proliferation, we observed a significant reduction in CAF abundance in the invasive margin and a concurrent decrease in the number of tumor cell strand structures formed over the 5-day GLAnCE assay (Figure 5E). This further suggested that modulating CAF supply in the invasive margin by curbing CAF proliferation was capable of diminishing tumor cell invasiveness in the GLAnCE system. Interestingly however, while CAF abundance in the invasive margin plateaued at the highest CAF densities tested (as seen in Figure 1D), the number of cancer cell strand structures continued to increase at these highest CAF densities (Figure 5D). This potentially suggests that changes in CAF abundance are not the only mechanism by which inhibition of EHMT2 in CAFs produces changes in tumor cell invasiveness and that likely a combination of both reduced CAF abundance and altered CAF EMT-inducing properties (e.g. secretion of cytokines, as observed in our GSA analysis) resulted in the reduced capacity of EHMT2-inhibited CAFs to induce tumor cell invasiveness.

## Discussion

CAFs play a central role in accelerating tumor growth and invasion via various direct and indirect effects on tumor cells including the upregulated secretion of growth factors, cytokines and chemokines that drive tumor cell growth and invasiveness, and the synthesis, deposition, and degradation of ECM components for architecture remodeling. Targeting CAFs to reduce these disease accelerating effects offers a potential strategy to augment standard of care treatment protocols. This work exploited a fully human *in vitro* culture model to identify the epigenetic enzyme EHMT2 in HNSCC CAFs as a target capable of decreasing both, CAF abundance within the tumor invasive margin and subsequent CAF-induced tumor cell invasiveness. RNA-seq analysis showed that inhibition of EHMT2 reduced secretion of EMT-inducing factors including POSTN, CXCL12, PTHLH and PTX3, and downregulated ECM-remodeling pathways suggesting a shift in CAF state towards a less activated phenotype with reduced potential to induce cancer cell invasiveness. Our RNA-seq analysis and preliminary SYN-1 staining results also suggest that EHMT2 inhibition potentially shifts CAF identity towards a neuronal-like phenotype through the de-repression of REST, thus dampening activated CAF-related functions. Functional data also suggest that inhibition of EHMT2 reduced CAF hyperproliferation, and to a lesser extent CAF migration.

Our observations are consistent with previously reported roles for EHMT2 such as mediating cell proliferation in cancer cells(*79, 82, 119–121*), myoblasts(*112, 113, 122*) and fibroblasts in the context of idiopathic pulmonary fibrosis(*49, 123, 124*). Our work therefore provides evidence that epigenetic regulation via EHMT2 can drive hyperproliferation in fibroblasts beyond the context of fibrosis, but also in CAFs in the context of the TME. Further EHMT2 function has been associated with the regulation of fibroblast activation state in the context of renal fibrosis(*80*) and with regulation of TGF-β1 driven changes in CAF CXCL9/CXCL10 expression during chemo-immunotherapy in pancreatic and colorectal cancers(*81*). Our findings are also in line with previous reports that inhibition of EHMT2 drives changes in CAF state such as changes in secretion profiles(*81*) and reprogramming of fibroblasts into alternative states(*108*).

Given the sensitivity of tumor cell invasive strand formation to CAF abundance, we speculate that the impact of EHMT2 inhibition on CAF proliferation and hence abundance may be the more dominant mechanism driving the observed reduction in tumor cell invasion. Increases in CAF abundance therefore could potentially amplify the effect of any CAF state changes. We thus propose a model, depicted in **Figure 6**, in which EHMT2 inhibition alters multiple aspects of the CAF phenotype resulting in a reduction in CAF abundance within the invasive margin via reduced CAF migration and proliferation, and subsequently a reduction in both the number and the capacity of CAFs present to induce tumor cell invasiveness. Overall, our findings support a model in which inhibition of EHMT2 in CAFs specifically deregulates the epigenetic determinants that constrain CAF to their pro-tumorigenic phenotype.

**Figure 6.**
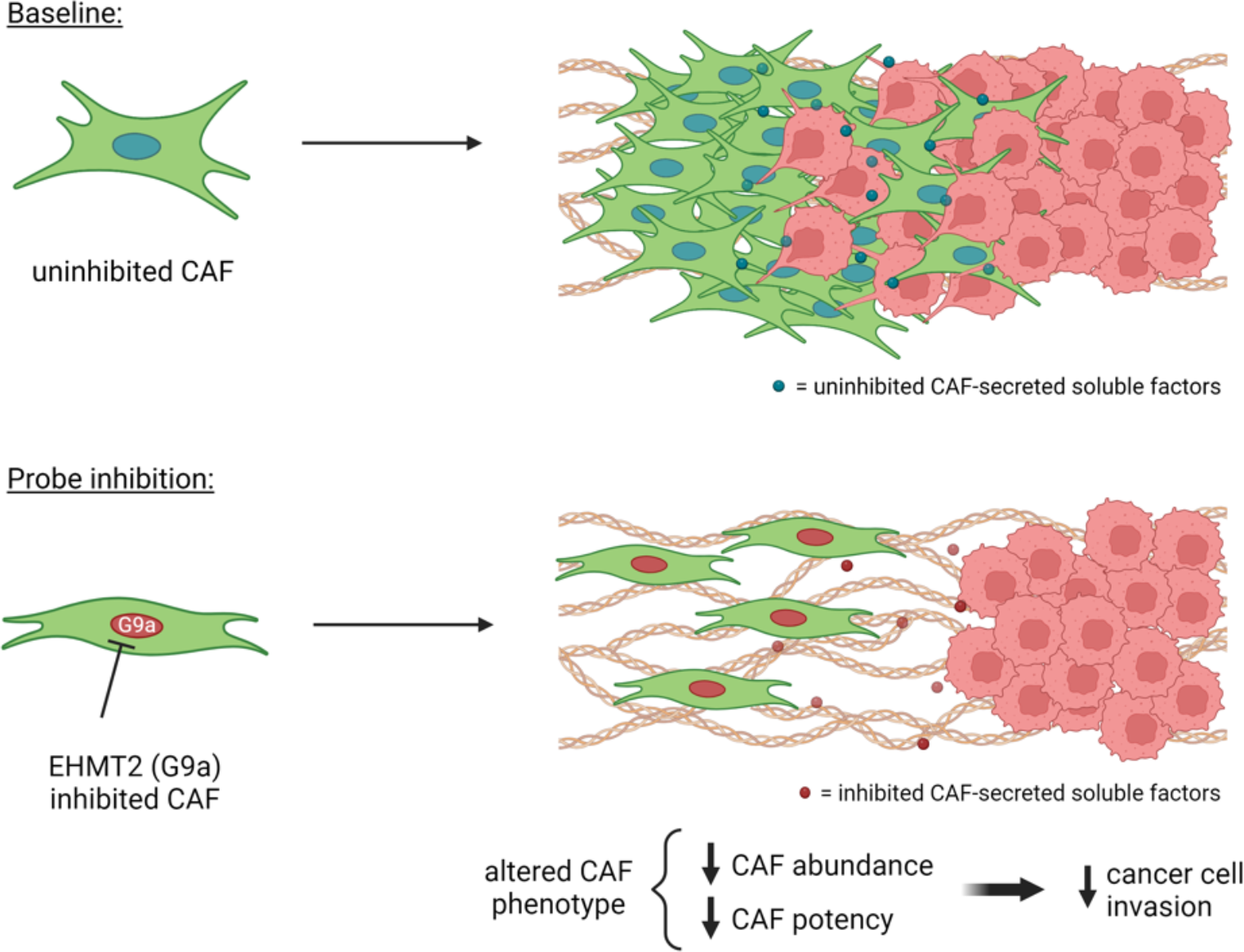
Proposed model of EHMT2 inhibition on CAF phenotype. Inhibiting epigenetic enzyme EHMT2 reduces CAF proliferation (and hence abundance in the invasive margin) and concurrently shifts HNSCC CAFs away from an activated, pro-tumorigenic phenotype, compared to baseline conditions. A reduction in CAF abundance compounds the impact of CAF de-activation resulting in significantly reduced CAF-induced cancer cell invasive strand formation.

High intra-tumor CAF content has been correlated with worse prognosis in HNSCC(*6, 7, 28*). Reducing CAF content by deletion of the entire CAF population has not shown clinical benefit and in fact resulted in increased tumor aggressiveness, at least in the context of pancreatic cancer(*125, 126*). Here we propose that epigenetically targeting CAF state to reduce both proliferation and activation, may provide an alternative strategy to reduce both CAF abundance and CAF “potency” within the tumor, while eliminating the risk of depleting anti-tumorigenic CAF populations. Our data provide additional evidence that EHMT2 is an epigenetic target with the potential to re-wire CAF phenotype, to both curb CAF proliferation and CAF-driven tumor invasiveness. The pro-invasive effect of systemic G9a inhibition in mouse models of tumor growth and metastasis may further, at least partially, be attributed to the effect of G9a inhibition in CAFs(*84, 85*). We note however, that previously reported direct inhibitors of G9a activity, including EZM-8266(*127*), were discontinued due to poor *in vivo* toxicity profiles(*82*), and EHMT2 appears to be an important regulator of proliferation and phenotype in various cell types.

Therefore, using CAF-targeting approaches or alternative molecules in the EHMT2 regulation pathway may represent more clinically tractable targets. In this study, we exploited a 3D *in vitro* platform, GLAnCE, to identify epigenetic modulators that regulate the activation state of head and neck squamous cell carcinoma (HNSCC) CAFs and subsequently their capacity to populate the tumor invasive margin and induce tumor cell invasiveness. Specifically, we found that inhibiting the epigenetic enzyme EHMT2 from methylating H3K9, re-wires CAF state to diminish both the supply of CAFs in the invasive margin zone and the capacity of the infiltrated CAFs to induce tumor cell invasiveness. The compounding effects of reduced CAF abundance via regulation of hyperproliferation, and reduced CAF “potency”, due to changes in CAF state, produced a significant reduction in CAF-induced tumor cell invasiveness. Overall, our data suggests that beyond fibrosis applications, EHMT2 is an interesting CAF-targeting candidate for pre-clinical follow-up. Further, our work showcases the potential of targeting epigenetic regulation of CAFs to induce a less tumorigenic phenotype as a strategy to reduce cancer cell invasiveness.

## Materials and Methods

### Cell Culture

The head and neck squamous cell carcinoma (HNSCC) cell line CAL33 (specifically oral squamous cell carcinoma) was purchased from DSMZ (Leibniz-Institute DSMZ, Germany) and mCherry-expressing CAL33 cells were generated using lentivirus transduction. Primary human oral squamous cell carcinoma CAFs (specifically tongue squamous cell carcinoma) were a gift from Dr. Ailles’ lab and isolated as previously described(*53*). CAFs were verified by negative staining for cytokeratin, CD45 and CD31, and positive staining for vimentin, and were used up to passage 12. GFP-expressing CAFs were generated using lentivirus transduction. HNSCC CAFs used in this study were derived from a single donor. Both cell types were maintained in IMDM (Wisent Bioproducts) supplemented with 10% FBS and 1% penicillin/streptomycin solution in plastic culture flasks, at 377C in 5% CO2. All cells were confirmed to be mycoplasma negative.

### Fabrication of Custom Device Components (for GLAnCE)

GLAnCE components were fabricated as previously reported(*52*). Briefly, GLAnCE polystyrene custom device bottoms were generated by hot embossing, using an array of 4 × 6 channel-shaped depressions with a depth of 276 µm ± 31 µm and distributed according to the well spacing of a 24-well plate, that was micro-milled into the surface of an aluminium stamp (*6061-T6, McMaster-Carr, CA).* Using hot embossing, the stamp was pushed into virgin transparent polystyrene (PS) (ST313120, GoodFellow) with 1700 N of force at 1907C for 6 minutes, with a temperature ramp and de-ramp of 50C/minute. Any excess melted polystyrene was removed using a heat-knife, to ensure the embossed PS sheet fit the dimensions of a black, bottomless 24-well plate (Greiner Bio-One). The channel-shaped depressions in the PS were termed “open channels” and were exposed to UV light at 254 nm and 15 Watts for 150 min to enhance collagen attachment(*54*).

To fabricate the polydimethylsiloxane (PDMS) slab used for generating GLAnCE gels, SYLGARD 184 Silicone Elastomer Kit was used according to the manufacturer’s recommended base to curing agent ratio (10:1) to a total weight of 30 g (27 g base + 3 g curing agent) (Sylgard 184, Ellsworth Adhesives). The mixture was degassed for 30 min in a vacuum chamber, poured into a 9 cm × 13 cm rectangular plastic mold, and baked at 100 ◦C overnight to cure. The PDMS slabs were then cut and removed from the plastic mold and interlaid between two laser-cut PMMA stencils containing 2 inlet and 2 outlet holes per channel, and the layered stencils and PDMS slab were clipped together to ensure proper alignment. Holes were then punched out through the stencil using a 1 mm biopunch (Integra). To prepare an adhesive layer to attach the cell-filled PS chip bottom to the black no-bottom 24-well plate, a sheet of double-sided, non-toxic adhesive tape (ARcare, Adhesive Research) was cut using a Silhouette Cameo Electronic Cutting Machine (Silhouette).

### Device Seeding & Assembly

The PS chip and PDMS slab were sandwiched together, to align each PS channel with 4 PDMS through-holes, thereby creating temporarily sealed channel structures with 2 inlets for hydrogel injection and 2 outlets for air escape, termed “closed channels”(*52*). These channels were then loaded with cell-gel suspension in a stepwise procedure: Firstly, 2.7 µL of mCherry+ cancer cells re-suspended in rat tail type I collagen hydrogel (ibidi) was injected through one side of the channel, whereby the volume injected dictated the length of the channel occupied by hydrogel. Next, the first hydrogel compartment containing mCherry+ cells was allowed to solidify for 5 min at 37°C and subsequently 3.4 µL of GFP+ CAFs re-suspended in rat tail type I collagen hydrogel were injected through the second side of the channel. Any air trapped between the two hydrogel fronts could escape through the lateral outlet holes. After this second injection the compartmentalized hydrogels were incubated at 37°C for 20 min to allow gelation of the second hydrogel compartment. Finally, the PDMS slab was peeled off, the PS chip with the channel-filled array of hydrogels was attached to the bottom of a black no-bottom, 24-well plate (Greiner Bio-One) using a cut double-sided, polyacrylic adhesive tape (Arcare, Adhesive Research), and cell culture media was immediately added to each well for further culture.

The selected seeding density for CAFs was 1 × 10^4^ cells/gel, to achieve an appropriate dynamic range of CAF infiltration in treatment conditions. For cancer cells, the seeding density selected was 0.5 × 10^4^ cells/gel, as this resulted in a cell density at day 5 suitable for the identification of strand structures. CAFs located at the hydrogel periphery contacted the PS channel walls and readily attached to both the PS channel walls and the collagen matrix, thereby preventing hydrogel peel-off from the channel wall over time; however an upper limit for CAF numbers was reached at 10^5^ CAFs seeded/hydrogel, beyond which hydrogels were observed to peel out of the channels. All experiments were therefore performed below this CAF seeding concentration.

### GLAnCE Invasion Assay

We performed GLAnCE co-culture invasion assays over 5 days, with GLAnCE plates being imaged on day 0 and day 5. Tumor invasion, fibroblast infiltration, and CAF growth were quantified on day 5. Tumor strand number was used as a surrogate measure of cancer cell invasiveness and were quantified by manually counting the number of strand structures per sample. A strand was defined as a cohesive structure of cancer cells clearly emerging from a cell aggregate, either connecting with another aggregate or protruding into the 3D hydrogel space. Fibroblast infiltration was assessed by quantifying the number of fibroblasts present beyond the initial CAF-tumor compartment interface using GFP signal intensity. An image from a media-only GLAnCE well was subtracted from all other images, to remove background GFP signal. To quantify changes in the number of fibroblasts in the invasive zone (a measure of CAF infiltration), Day 0 and day 5 images were overlaid, and a 1000 µm × 1000 µm region-of-interest (ROI) was positioned to the right and immediately adjacent to the day 0 GFP/mCherry interface. To estimate fibroblast occupancy within this defined ROI, the day 5 GFP signal was then quantified by thresholding and calculating the % of non-black pixels in the ROI masked image. For each replicate, the signal threshold level to define GFP+ pixels was defined manually and kept consistent throughout the quantification of all images within that specific replicate. To quantify CAF growth, a ROI was defined in the CAF compartment, excluding the invasive margin, for each sample. This was done to account for minor variations in the positioning of the CAF and cancer cell interface across samples. The MGV of the GFP+ signal of the ROI was measured on D0 and D5, then the value on D5 was normalized to the corresponding channel’s D0 MGV measurement.

### Atomic Force Microscopy & Scanning Electron Microscopy

To assess the structure of collagen hydrogels with or without cells, we used scanning electron microscopy (SEM). Specifically, rat tail type I collagen hydrogels with or without encapsulated cells were cultured for 5 days and then decellularized using puromycin treatment at 5 µg/mL for 48 hrs, followed by a 48 hr PBS wash on a shaker with a change of PBS at 24 hr. Decellularized gels were then dehydrated in 100% ethanol (anhydrous) overnight. After dehydration, the gels were critically point dried (CPD) in a Autosamdri-810 (Tousimis, USA) with a purge phase of 20 minutes. After CPD, gels were mounted onto SEM chucks that had been coated with carbon tape and samples were gold-palladium sputter coated for 60 secs (Polaron SC7640 Sputter Coater; Quorum Technologies, Canada). To image the gels, scanning electron microscopy (Hitachi S-3400N, Hitachi High Technologies Canada) at 10 kV was utilized.

To assess the mechanical properties of GLAnCE gels in different cellular compartments, we used Atomic Force Microscopy (AFM), as performed as previously described(*55*). GLAnCE gels with different seeding conditions were tested using a commercial AFM (Bioscope Catalyst, Bruker, Santa Barbara, CA) mounted on an inverted optical microscope (Nikon Eclipse-Ti). Force mapping was accomplished using a pyramidal tip and 3-4 regions were measured per gel with multiple readings collected per region, with a 10 nN consistent trigger force applied to the substrate. All AFM measurements were collected at room temperature.

### Epigenetic Inhibitor (Probe) Library Screen

6 × 10^4^ CAFs were plated per well in a 6-well plate and probes were added at a concentration of 1 µM in complete medium on day 1 and replaced with fresh probes in complete medium on day 5. On day 7, CAFs were harvested by trypsinization, pelleted and cell numbers were counted. For each probe-treated CAF population, 1.5 × 10^4^ CAFs were then resuspended in rat tail type 1 collagen and seeded into the second compartment of GLAnCE gels adjacent to HNSCC cancer cells in the first compartment. Cultures were then incubated for a 5-day compartmentalized co-culture before quantification of CAF invasion and tumor cell strand formation. Experiments were performed in triplicate and all metrics were normalized to DMSO-treated CAF co-cultures.

### Cell Viability Assay

To assess cell viability in cultures, live-dead staining was performed using Calcein AM (20 632, Cayman Chemical Co.) at 1:1000 and propidium iodide (PI) (ThermoFisher Scientific) at 1:50. Prior to adding reagents, samples were washed once with PBS to remove media remnants, then incubated with reagents for 30 min at room temperature, away from light. Samples were washed twice for 5 min each with PBS before imaging. Stains were then assessed using fluorescence microscopy (see below). PI intensity was normalized to cell number in the field of view analyzed.

### Immunofluorescent Assessment of Probe Treated Cells

Glass coverslips were placed in 12-well plates, UV sterilized for 15 minutes, then coated with poly-d-lysine for 1 hour at 37C. After aspirating poly-d-lysine solution and allowing coverslips to dry, CAFs were seeded at 15 000 cells/well. Probes or DMSO (vehicle) were added on day 1 and day 5 (identical to screen protocol). For H3K9me2 staining, parallel coverslips were fixed on days 2, 6, 8 and 10 with 4% paraformaldehyde (PFA) for 10 minutes at room temperature, then washed three times with PBS. Upon the collection of all timepoints, the following was performed: coverslips were blocked using 1X PBS/5% BSA/0.3% Triton X-100 for 60 minutes at room temperature, then incubated with primary mouse anti-dimethylK9H3 antibody (abcam ab1220) diluted to 1:200 in dilution buffer (1X PBS/1% BSA/0.3% Triton X-100) for 2 hours at room temperature. Coverslips were washed three times with PBS prior to incubating with secondary chicken anti-mouse AlexaFluor 647 antibody (Thermo Fisher Scientific A21463) diluted to 1:200 in dilution buffer for 1.5 hours at room temperature, away from light, and washed again three times in PBS. To stain nuclei, nuclear stain Hoechst (20 mM; Thermo Fisher Scientific) was added at 1:1000 dilution for 15 minutes at room temperature. Finally, coverslips were washed one final time prior to mounting onto a microscope slide and sealing, then imaged with a confocal microscope. A similar workflow was performed with primary anti-SYN-1 antibody (Abcam, ab8) diluted to 1:2000 in 3% normal donkey serum (Millipore Sigma, S30) and 0.3% Triton-X, secondary donkey anti-rabbit AlexaFluor 488 antibody (Thermo Fisher Scientific) diluted to 1:1000, followed by nuclear staining using 300 nM of DAPI (Thermo Fisher Scientific, D3571). For BODIPY staining, 7 days after seeding CAFs onto poly-d-lysine coated coverslips, samples were fixed with 4% PFA for 10 minutes at room temperature. A master mix of BODIPY (ThermoFisher Scientific) diluted to 1:1000 and DRAQ5 (eBioscience) diluted to 1:1000 in PBS was prepared and incubated on samples for 30 minutes at room temperature, away from light. Coverslips were washed three times with PBS, mounted onto a microscope slide and sealed, then imaged using confocal microscopy.

### Quantification of Co-Culture Growth

To assess the impact of probe treatment on tumor cell growth, we performed a co-culture growth assay, in which HNSCC cancer cell line and CAF cells were re-suspended together in the rat tail type I collagen and seeded into GLAnCE with one injection of 5.7 µL to generate a single compartment, fully co-mixed gel. Images were acquired every day over a period of 5 days, and growth of cancer cells specifically was quantified using the mean gray value of the mCherry channel.

### Quantification of Probe Treated CAF Migration

A Transwell® migration assay was utilized to quantify the migratory potential of probe-treated CAFs compared to the baseline phenotype. For the Transwell® migration assay, 6.5 mm Transwells® with 8.0 µm pore polyester membrane inserts were used. After 7 days of treatment with probe or DMSO, CAFs were dissociated with Trypsin and counted. 100 µL of serum-free IMDM was first added to the top chamber, followed by 200 µL of 2.5 × 10^5^ CAFs/mL in serum-free IMDM. Finally, 750 µL of complete IMDM was added to the lower chamber. Transwells® were then incubated for 10 hours at 377C in 5% CO2, then washed, fixed with 4% PFA, washed again, and then stained with Crystal Violet. Transwells® were washed one final time and a cotton swab was used to scrape off the non-invasive CAFs (inside the chamber), before placing on a microscope slide to be imaged using light microscopy. The intensity of the Crystal Violet in images of the stained cells was used to quantify the number of cells that invaded through the Transwell® membrane relative to a control with untreated CAFs.

### Quantification of Probe-treated CAF Contractility

After 7 days of probe or DMSO treatment of CAFs, cells were detached and counted. Rat tail type I collagen gel plugs were then prepared by re-suspending 100 000 CAFs/500 µL of hydrogel in a 24-well plate well. Gel plugs were placed in the incubator for 30 minutes to solidify, then serum-free IMDM was added to each well. Gel plugs were immediately detached from the side of the well walls by running a pipette tip around the circumference of the well, taking care not to rip the gel plug. Gel plugs were cultured for 5 days, during which time the hydrogel contracted to varying extents dependent on the contractility of the encapsulated cells. Gels were imaged on day 5 to enable quantification of the % change in gel area between day 0 and 5.

### Bulk RNA-sequencing Sample Preparation

Cells were treated with probe or DMSO for 7 days, before RNA extraction using the Norgen Biotek Single Cell RNA Purification Kit (Norgen Biotek Corp. #51800). RNA samples were then quantified by qubit (Life Technologies) and an Agilent Bioanalyzer was used to assess RNA quality. All samples had a RIN above 8. A SMART-Seq v4 Ultra Low Input RNA Kit for Sequencing (Clontech #634894) was used per manufacturer’s instructions for amplification of RNA and subsequent cDNA synthesis. AMPure XP Bead (Agencourt AMPure beads XP PCR, Beckman Coulter A63881) purification was done manually for the first amplification set. A bead ratio of 1x was used (50 µL of AMPure XP beads to 50 µL cDNA PCR product with 1 µL of 10x lysis buffer added, as per Clontech instructions), and purified cDNA was eluted in 17 µL elution buffer provided by Clontech. All samples were quantitated using Bioanalyzer 2100 Instruments (Agilent Genomics). All samples proceeded through NexteraXT DNA Library Preparation (Illumina FC-131-1096) using NexteraXT Index Kit V1 or V2 Set A (FC-131-1002 or FC-131-2001) following the manufacturer’s instructions. An aliquot of all samples was first normalized to 150 pg/µL with Nuclease-Free Water (Ambion), then this normalized sample aliquot was used as input material into the NexteraXT DNA Library Prep. AMPure XP bead purification was done using 0.9x bead ratio to sample volume, and all samples were eluted in 22 µL of Resuspension Buffer (Illumina). As with the Amplification sets, manual bead purification was done for the first Library set. All samples were run on Agilent Bioanalyzer 2100 using High Sensitivity DNA chips. A portion of this library pool was sent to an outside vendor for sequencing on an Illumina NextSeq HighOutput single read. An average of 400M reads were obtained per pool, with an average of 40M reads/sample across the entire data set.

## Bulk RNA-sequencing Analysis

### Preprocessing

The base calls were demultiplexed and converted to FASTQ format using Illumina bcl2fastq tool. Quality control analysis was done with FastQC and MultiQC(*56*). Raw FASTQ sequencing reads were trimmed using the Trimmomatic tool with the default parameters to remove reads with adaptor contamination or poor quality(*57*). The trimmed reads were then aligned to the GRCh38.v84 genome and dbSNP.v144 database using HISAT v2.1.0(*58*) to generate aligned reads. Alignments were compressed and sorted using SAMtools(*59*). featureCounts v1.6.3(*60*) tool was used to quantify expression by counting the number of reads that map to genes for all the samples.

### Differential Expression Analysis

To perform principal component analysis, we first transformed normalized counts using the variance stabilize transformation implemented in DESEq2(*61*). The vst normalized count data was then z-scaled for visualization purposes.

Differential gene expression was done using raw counts data inputted into the DESeq2 R package. We removed all genes that had 10 or fewer read count in sum across all samples. With the filtered expression matrix, we set the DMSO condition as the basal state and calculated the shrunken log fold changes between genes using the apeglm R package(*62*) and tested for significance using the Wald test implemented in DESeq2.

### GSEA Analysis

To calculate gene set enrichment across the MSigDB database, we used the GSEA function implemented in the clusterProfiler R package(*63*) in tandem with the msigdb R package(*64*). We aggregated the p-values between the OICR and A366 iterations using the Fisher’s-method implement in the metap R package(*65*), and adjusted for multiple hypothesis testing using an FDR correction.

### TF Analysis

The ChEA3(*66*) libraries ARCHS4_Coexpression (ARCHS4), ENCODE_ChIP-seq (ENCODE), Enrichr_Queries, GTEx_Coexpression (GTEx), Literature_ChIP-seq (Literature), and ReMAP_ChIP-seq (ReMAP) were downloaded from https://maayanlab.cloud/chea3/ in April-2022; their corresponding md5sum identifiers can be found in the git repository associated with this analysis. Using the GSEA function from the R package clusterProfiler(*67*), we calculated the p-values and normalized enrichment score (NES) for every geneset in the ChEA3 databases. We then aggregated the p-values between the OICR and A366 iterations using the Fisher’s-method implement in the metap R package(*65*), and adjusted for multiple hypothesis testing using an FDR correction. For visualization, we then reduced the NES per dataset to the mean score across all replicates for each transcription factor. From this averaged-NES, we selected the top 5 TFs with the smallest q-values in each direction.

### Data & Code Availability

The code used to process the aligned data is found at https://github.com/mcgahalab/hnscc_cafs.

### Microscopy & Image Analysis

All widefield images were acquired at 4X using an ImageXpress Micro or ImageXpress Pico high-content system (Molecular Devices, USA), using both laser-based and image-based autofocus settings. Confocal images of live or fixed samples were acquired using a Leica SP8 confocal microscope (Leica Microsystems, Germany). The former was required when measuring gel height of seeded gels on day 0 and day 5. Image analysis was performed using FIJI(*68*) with custom macros.

### Statistical Analysis

All results are presented as mean ± standard error of the mean (SEM) from multiple independent experiments, with at least 3 biological replicates, unless mentioned otherwise. All data was analyzed using GraphPad Prism 6 (GraphPad Software, La Jolla, Ca, USA). Two test groups were compared using an F-test to determine equal variance. An unpaired t-test was performed whenever equal variance could be assumed, and an unpaired t-test with Welch’s correction whenever unequal variances were assumed. For 3 test groups or more, a one-way analysis of variance (ANOVA) was performed, with Dunnett’s multiple comparisons to compare between groups. All tests were two-tailed, and asterisks represent the following p-value cut-offs: * < 0.05, ** < 0.005, *** < 0.0005, **** < 0.0001.

## Data and materials availability

All data needed to evaluate the conclusions in the paper are present in the paper and/or the Supplementary Materials.

