## Supplemental Information for "EHMT2/G9a-Inhibition Reprograms Cancer-Associated Fibroblasts (CAFs) to a More Differentiated, Less Proliferative and Invasive State"

Nila C Wu *et al.*

**This PDF file includes:**

Figs. S1 to S2
Table S1

**Fig. S1.**

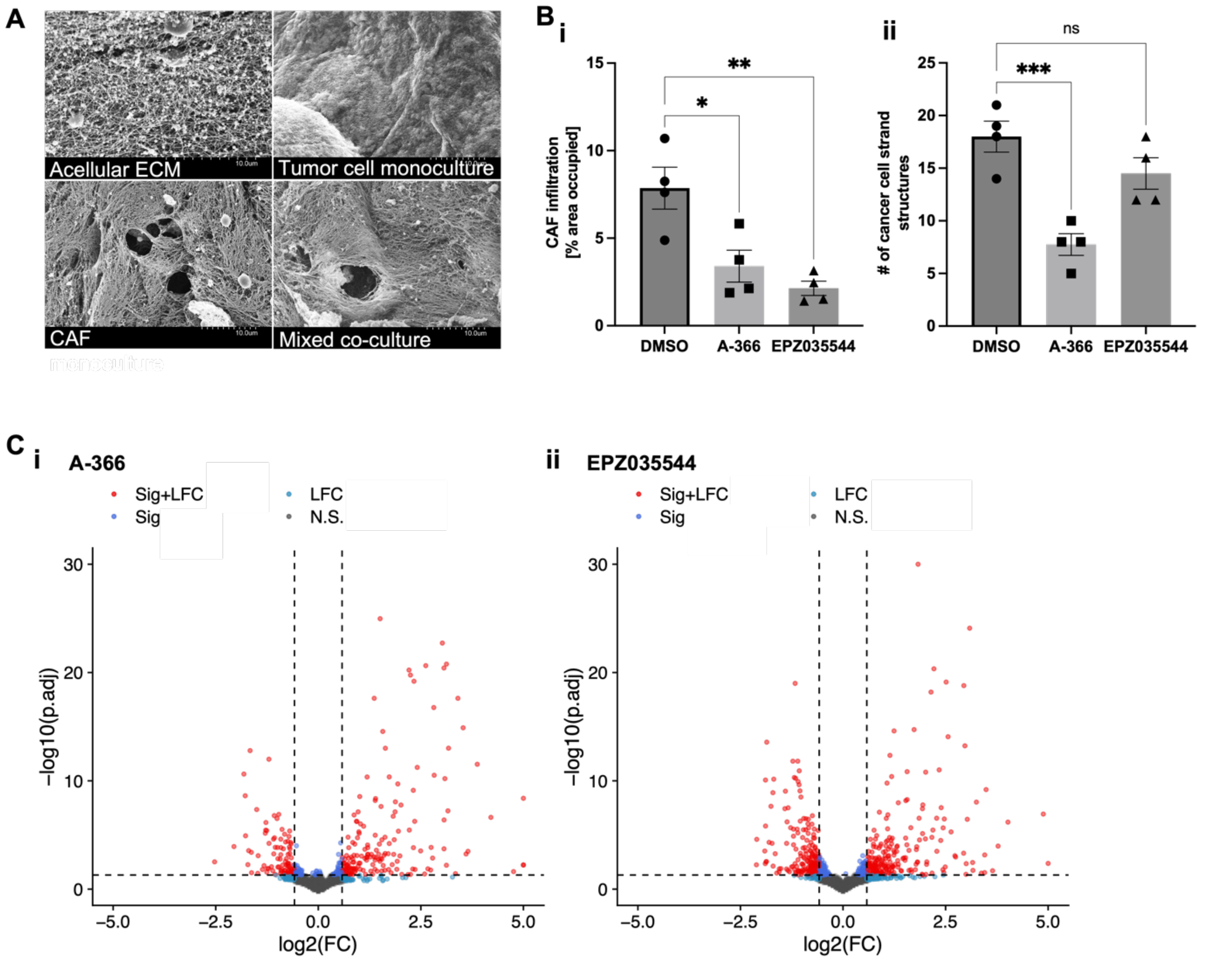


**SI Figure 1. (A)** CAF-mediated ECM remodeling results in distinctive pores within the ECM visible by SEM. These pores are only observed in the presence of CAFs, in either mono-culture of mixed co-culture. **(B)** Compartmentalized GLAnCE co-cultures seeded with EHMT2/G9a-inhibited CAFs adjacent to a different HNSCC cancer cell line, mCherry-transduced CAL27s, similarly results in a reduction of **(i)** CAF infiltration and **(ii)** number of cancer cell strand structures. Results show mean ± SEM for n = 4 biological replicates. **(C)** Volcano plots obtained after bulk RNA-seq of **(i)** A-366 treated CAFs and **(ii)** EPZ035544 treated CAFs, compared to DMSO treated CAFs.

**Fig. S2.**

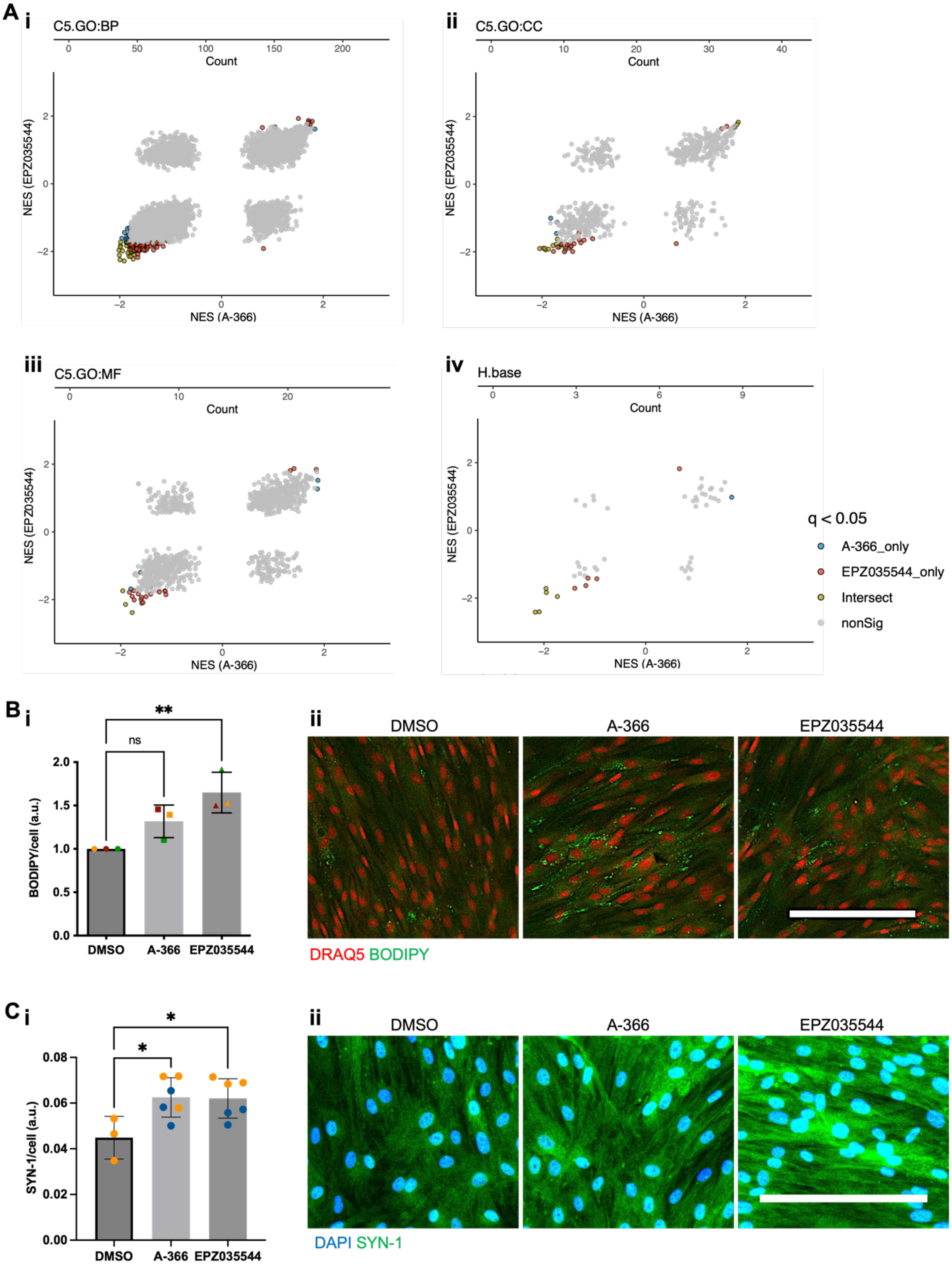

A
i
C5.GO:BP
Count
NES (EPZ035544)
NES (A-366)
ii
C5.GO:CC
iii
C5.GO:MF
iv
H.base
q < 0.05
A-366_only
EPZ035544_only
Intersect
nonSig
B
i
BODIPY/cell (a.u.)
ns
**
DMSO
A-366
EPZ035544
ii
DRAQ5 BODIPY
C
i
SYN-1/cell (a.u.)
*
ii
DAPI SYN-1

**SI Figure 2.** **(A)** Comparison of the enriched gene set directionality between A-366 and EPZ035544, of the 4 collections: **(i)** GO:BP, **(ii)** GO:CC, **(iii)** GO:MF, and **(iv)** Hallmark. Enriched gene sets between inhibitors mostly share the same directionality, with two pathways in the GO collections that had mismatched directionality between treatments. **(B)** BODIPY staining to assess lipid droplet content between CAF treatments. **(i)** No significant difference of BODIPY intensity was observed. **(ii)** Representative BODIPY stained images. Scale bar is 200 μm. Results show mean ± SEM for N = 4 biological replicates. **(C)** **(i)** SYN-1 staining to assess a neuronal marker. Note, N = 2 for A-366 and EPZ035544, and N = 1 for DMSO, therefore statistics were performed on all technical samples available; n = 6 technical replicates for A-366 and EPZ035544, n = 3 technical replicates for EPZ035544. **(ii)** Representative SYN-1 stained images. Scale bar is 200 μm.

**Table S1.**

| Compound Name | Target |
|---|---|
| A-196 | MT: SUV420H1/H2 |
| A-366 | MT: G9a/GLP |
| A-395 | Kme: EED |
| A-485 | AT: CBP, p300 |
| BAY-299 | BRD: BRPF2/TAF1 |
| BAY-6035 | MT: SMYD3 |
| BAY-598 | MT: SMYD2 |
| BAY-876 | BMP2K/BIKE |
| BAZ2-ICR | BRD: BAZ2A/2B |
| GSK-864 | DEHYDR: IDH1 mt |
| GSK-4027 | BRD: PCAF/GCN5 |
| GSK-484 | PAD: PADI4 |
| GSK-591 | MT: PRMT5 |
| GSK-8814 | BRD: ATAD2A/B |
| JQ1 | BRD: BET family |
| LLY-283 | MT: PRMT5 |
| MS023 | MT: PRMT type 1 |
| MS049 | MT: PMRT4,6 |
| NI-57 | BRD: BRPF1/2/3 |
| NVS-1 | BRD: CECR2 |
| OICR-9429 | WD40: WDR5 |
| PFI-2 | MT: SETD7 |
| PFI-3 | BRD: SMARCA2/4 |
| PFI-4 | BRD: BRPF1B |
| PFI-5 | MT: SMYD2 |
| SGC707 | MT: PRMT3 |
| SGC0946 | MT: DOT1L |

| | |
|---|---|
| SGCCBP30 | BRD: CREBBP/EP300 |
| TP-064 | MT: PRMT4 |
| TP-472 | BRD: BRD9/7 |
| UNC1215 | Kme: L3MBTL3 |
| UNC1999 | MT: EZH2/H1 |

**SI Table 1. List of epigenetic regulator probes in screened library, and corresponding targets.**
